# NICEgame: A workflow for annotating the knowledge gaps in metabolic reconstructions, using known and hypothetical reactions

**DOI:** 10.1101/2022.02.10.479881

**Authors:** Evangelia Vayena, Anush Chiappino-Pepe, Homa MohammadiPeyhani, Yannick Francioli, Noushin Hadadi, Meriç Ataman, Jasmin Hafner, Stavros Pavlou, Vassily Hatzimanikatis

## Abstract

Advances in medicine and biotechnology rely on the further understanding of biological processes. Despite the increasingly available types and amounts of omics data, significant biochemical knowledge gaps remain uncharacterized. Several approaches have been developed during the past years to identify missing metabolic annotations in genome-scale. However, these approaches suggest missing metabolic reactions within a limited set of already characterized metabolic capabilities. In this study, we introduce NICEgame (<u>N</u>etwork Integrated <u>C</u>omputational <u>E</u>xplorer for <u>G</u>ap <u>A</u>nnotation of <u>Me</u>tabolism), a workflow to characterize missing metabolic capabilities in genome-scale metabolic models using the ATLAS of Biochemistry. NICEgame suggests alternative sets of known and hypothetical reactions to resolve gaps in metabolic networks, assesses their thermodynamic feasibility, and suggests candidate genes and proteins to catalyze the introduced reactions. We use gene essentiality data use to identify metabolic gaps in the latest genome-scale model of *Escherichia coli*, iML1515. We apply our gap-filling approach and further enhance its genome annotation, by suggesting reactions and putative genes to resolve 46 % of the false negative gene essentiality predictions.

## 1. Introduction

Fully defined metabolic networks and annotated genomes can provide a holistic picture of the cell function and enable the design of more robust and effective bioengineering and drug targeting strategies. However, all known replicating genomes miss functional annotations for a fraction of the open reading frames. For example, the best characterized organism, i.e., *Escherichia coli*, lacks annotation for approximately 1600 genes, which represents 35% of its total number of genes^1^. A limited knowledge of the cell function is especially troublesome in infectious pathogens and organisms that could be used as a chassis in the industry to produce valuable compounds. Systematically identifying missing knowledge of metabolic capabilities of the cell and accelerating the functional annotation of genomes can save lives, time, and costs involved in the design of medical therapies and biotechnology projects.

Systematic analysis of metabolic functions and identification of knowledge gaps relies on the gold standard of computational models of metabolism. Genome-scale models are organism-specific databases of all known metabolic functions. They have been widely used to study the metabolism of model organisms, such as *E. coli*^2^ and yeast^3^, and pathogens such as *S. typhimurium*^4^ and *P. falciparum*^5^, to identify host-pathogen interactions^6^, drug targets^7^, and metabolic engineering strategies^8^, among others^9^. Genome-scale models rely on the functional annotation of genes for their reconstruction. High-quality gene annotation leads to a proper prediction of the cellular physiology. Genome-scale models, hence, represent a powerful framework to identify missing knowledge through false predictions.

Approaches to perform functional annotation of genomes involve experimental^10^ (e.g. in vitro assays) and bioinformatics^11^ (e.g. blasting) methods. However, experiments require specific hypotheses and are time-consuming and blasting and other computational approaches have been limited to the space of known annotated proteins and biochemistry.

Exploring the space on unknown biochemistry is required to accelerate our understanding of the cell function and include in cells novel chemistry. The strategies to explore such unknown biochemical space are primarily based on machine learning (ML) or mechanistic approaches^12,13^.

Recently, an ATLAS of Biochemistry was constructed based on a mechanistic understanding of the enzyme function^14^. The ATLAS of Biochemistry includes over 150,000 putative reactions between known metabolites. Hence, it represents the upper limit of the possible biochemical space and allows an efficient exploration of the uncharacterized metabolic functions in cells. Furthermore, the tool BridgIT^15^ was developed to identify putative genes and proteins catalyzing a reaction. The potential of a combined use of ATLAS of Biochemistry and BridgIT to identify unknown metabolic annotations is tremendous but remains unexplored.

In this study, we present a <u>N</u>etwork Integrated <u>C</u>omputational <u>E</u>xplorer for <u>G</u>ap <u>A</u>nnotation of <u>Me</u>tabolism (NICEgame). NICEgame combines the analysis of metabolic functions using genome-scale models (GEMs), exploration of unknown biochemistry using the ATLAS of Biochemistry, and identification of uncharacterized genes using BridgIT. We applied NICEgame to suggest novel biochemistry in *E. coli*’s strain MG1655 and further enhance its genome annotation. We identified metabolic gaps in *E. coli* responsible for 146 false gene essentiality predictions in glucose minimal media. We propose 77 biochemical reactions linked to 35 candidate genes to fill 46% of these gaps. We integrated this information into a thermodynamically curated genome-scale model of *E. coli* that we name iEcoMG1655. The iEcoMG1655 metabolic model increased the essentiality prediction accuracy by 23.5% with respect to its predecessor iML1515^2^. Finally, the NICEgame workflow is applicable to any organism or cell with a genome-scale model and is available as a GitHub repository (https://github.com/EPFL-LCSB/NICEgame), with the combined use of available online resources: The ATLAS of Biochemistry^14^ and BridgIT^15^. Overall, NICEgame is a workflow for rapid and systematic identification of metabolic gaps, missing biochemistry, and candidate catalyzing genes. Hence it will accelerate the complete identification of metabolic functions and annotation of genomes, and with it enable the design of robust bioengineering and drug targeting strategies.

## 2. Results and Discussion

### 2.1 A workflow to identify and curate gaps in metabolic networks

To identify metabolic gaps in the GEM of *E. coli*, we developed a Network Integrated Computational Explorer of Gap Annotation for Metabolism Expanded (NICEgame). NICEgame leverages the potential of the ATLAS of Biochemistry^14^ as a database of novel biochemistry, i.e. not yet observed reactions, and the optimization-based exploration of metabolic models to identify missing biochemistry. Moreover, we couple NICEgame with a method to map orphan biochemistry to genes called BridgIT^15^. In this way, NICEgame optimally identifies metabolic gaps, finds putative missing biochemistry, evaluates alternative solutions, and identifies the top suggested reaction and associated catalyzing enzyme and gene.

NICEgame involves seven main steps (Figure 1). The first step involves the harmonization of metabolite annotations with the ATLAS of Biochemistry. This is a necessary step to assure the proper connectivity of the metabolites in the GEM and the database. The second step comprises a preprocessing of the GEM (e.g., by defining the media). In the third step, NICEgame merges the GEM and ATLAS of Biochemistry. Thereafter, we call this merged network ATLAS-merged GEM. This process can follow various strategies as we will discuss below. The fourth step includes a comparative essentiality analysis with the isolated and ATLAS-merged GEM. At this point, we identify the reactions or genes that are *in silico* essential for a phenotype (by default growth) in the GEM and dispensable in the ATLAS-merged GEM. We define such reactions or genes as *rescued*. The rescued reactions and genes will be the targets for gap-filling. In the fifth step, NICEgame systematically identifies alternative biochemistry to the rescued reactions or genes. By default, NICEgame suggests a minimal number of reactions to be added to the GEM. In the sixth step, we evaluate and rank all alternative biochemistry. Last, NICEgame identifies a gene that can catalyze the top-ranked suggested biochemistry using the BridgIT tool.

**Figure 1.**
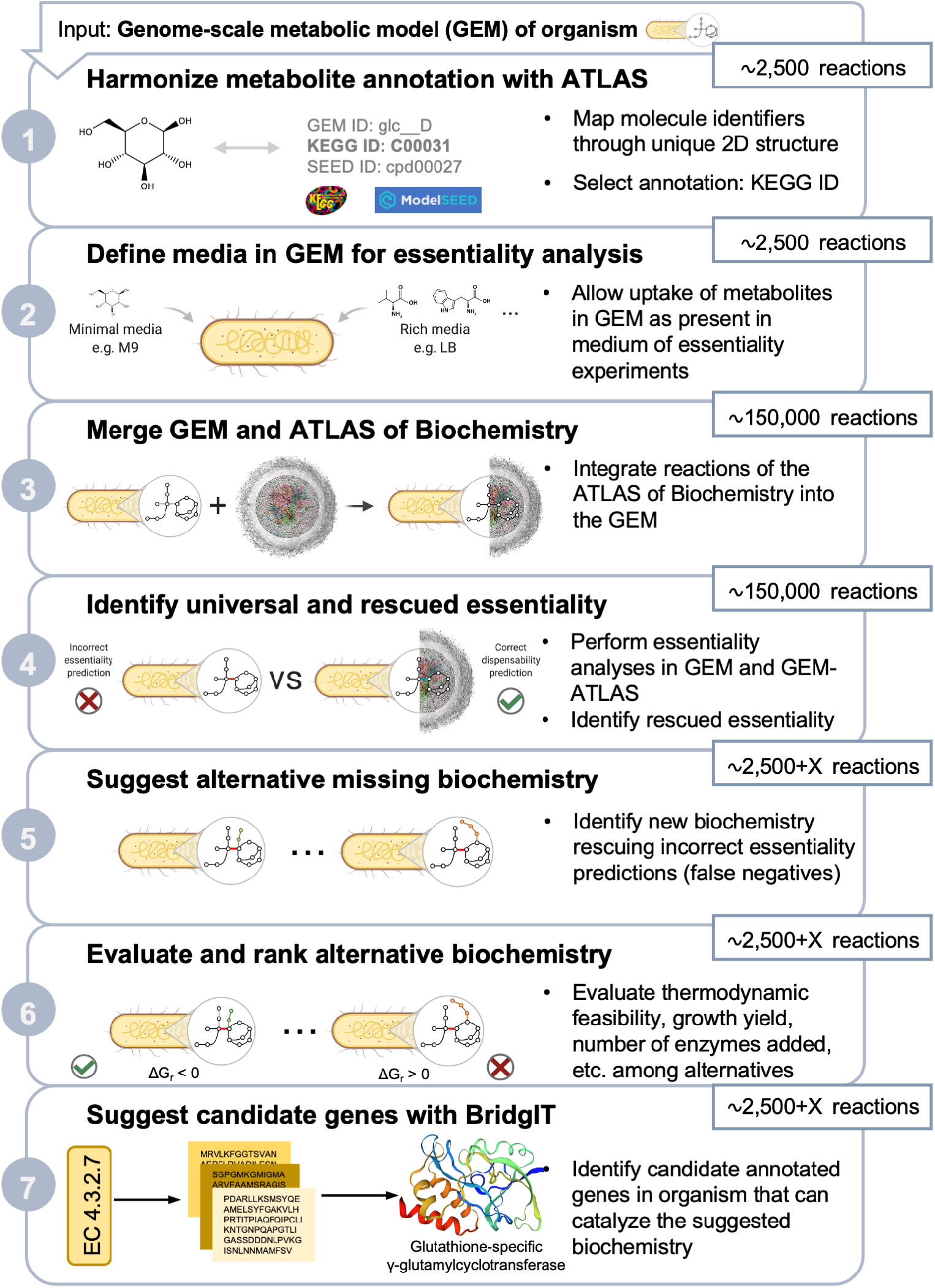
Pipeline to construct and use the NICEgame workflow to annotate missing metabolic functions. The NICEgame workflow uses a genome-scale metabolic model (GEM) as input. (1) We first harmonize the annotations of the GEM metabolites to map them to compounds in the ATLAS of Biochemistry. (2) We define the conditions for subsequent essentiality analyses, i.e., the media composition. (3) The original GEM is merged with ATLAS and (4) an essentiality analysis is performed in the original and the expanded network to identify which gaps can be rescued. (5) Alternative sets of biochemistry are generated to fill in the gaps and (6) are then evaluated. (7) At the last step, we use BridgIT to identify catalyzing for the suggested biochemistry.

The alternatives are evaluated based on the impact they have on the metabolic network and the performance of the model (Figure 2). Solution sets, i.e., sets of reactions that are added to the network to reconcile a gap, that result in a higher yield or do not affect the yield are preferred to solutions that reduce the flexibility of the model, whereas solutions that expand the metabolome or the enzymatic capabilities of the original model are ranked lower. Another criterion is the number of reactions that are used to complement each rescued reaction. Usually, organisms do not favor larger pathways since they normally require more protein production, which is a highly energetically demanding process. We assumed that short pathways would be preferred to perform a metabolic function. Lastly, the alternatives that increase the ability of the model to correctly reproduce knockout phenotypes and do not add redundancy are ranked higher. These criteria are converted into scores (See Methods). A positive value for any of the scores penalizes the alternative solution set in our ranking system.

**Figure 2.**
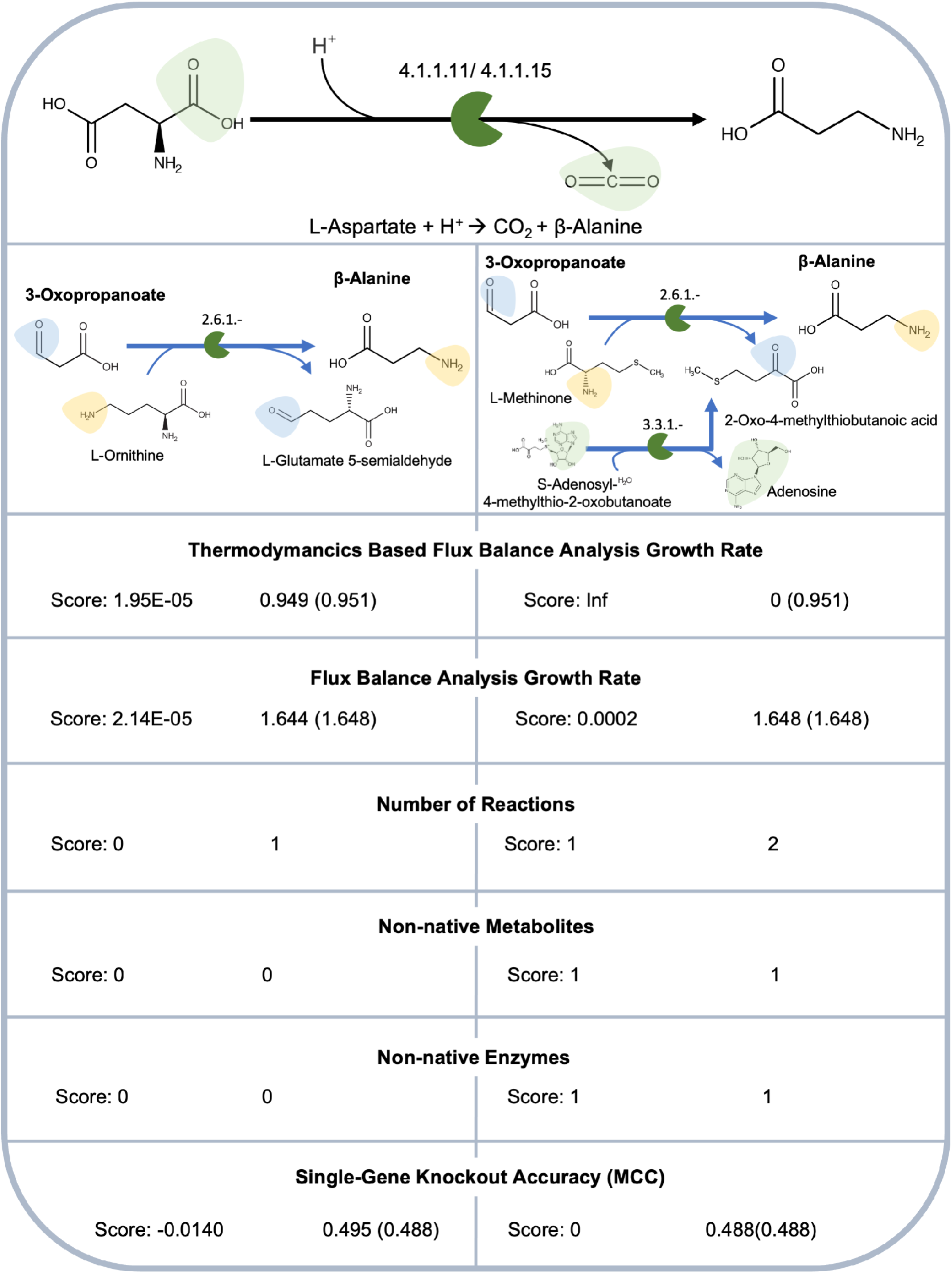
Rating of the alternative reaction sets. Two alternatives to reconcile the false negative prediction for the gene panD are shown. The second alternative is rejected because it is thermodynamically infeasible. This alternative is penalized in all scores (i.e., all scores are positive) since it adds two reactions, one non-native metabolite and one non-native enzymatic capability to the network. Regarding yield, the first alternative is also penalized (i.e., positive score) since it constrains the model both with and without thermodynamic constraints. However, it performs well at the reaction, metabolite and enzyme scores (i.e., they are all equal to 0) and it increases the accuracy of the model (i.e., MCC score is negative).

### 2.2 Identification of metabolic gaps in Escherichia coli with NICEgame

Our analysis identified 146 False Negative gene essentiality model predictions (Figure 3A) that translate into 152 false negative essential reactions (Supplementary Table S1). As a case study, we used two different subsets of ATLAS to gap-fill the metabolic network of iML1515. The first subset, Ecoli_mets_ATLAS DB, expands the reaction space of the model, by adding reactions involving only metabolites from the iML1515 reconstruction. The second subset, Ecoli_Yeast_mets_ATLAS DB, expands the reaction and metabolite space of the model, by adding reactions involving only metabolites from the iML1515 and Yeast8 metabolic reconstructions. In total, we could suggest at least one thermodynamically feasible solution set for 93 out of the 152 false negative reactions (Figure 3B). However, gaps, that cannot be resolved by our approach, still remain in the model. The falsely negative gene pabA (b3360) regulates the synthesis of 4-amino-4-deoxychorismate, a precursor of Folate, from Chorismate (Figure 3C), constitutes such an unresolved metabolic gap.

**Figure 3.**
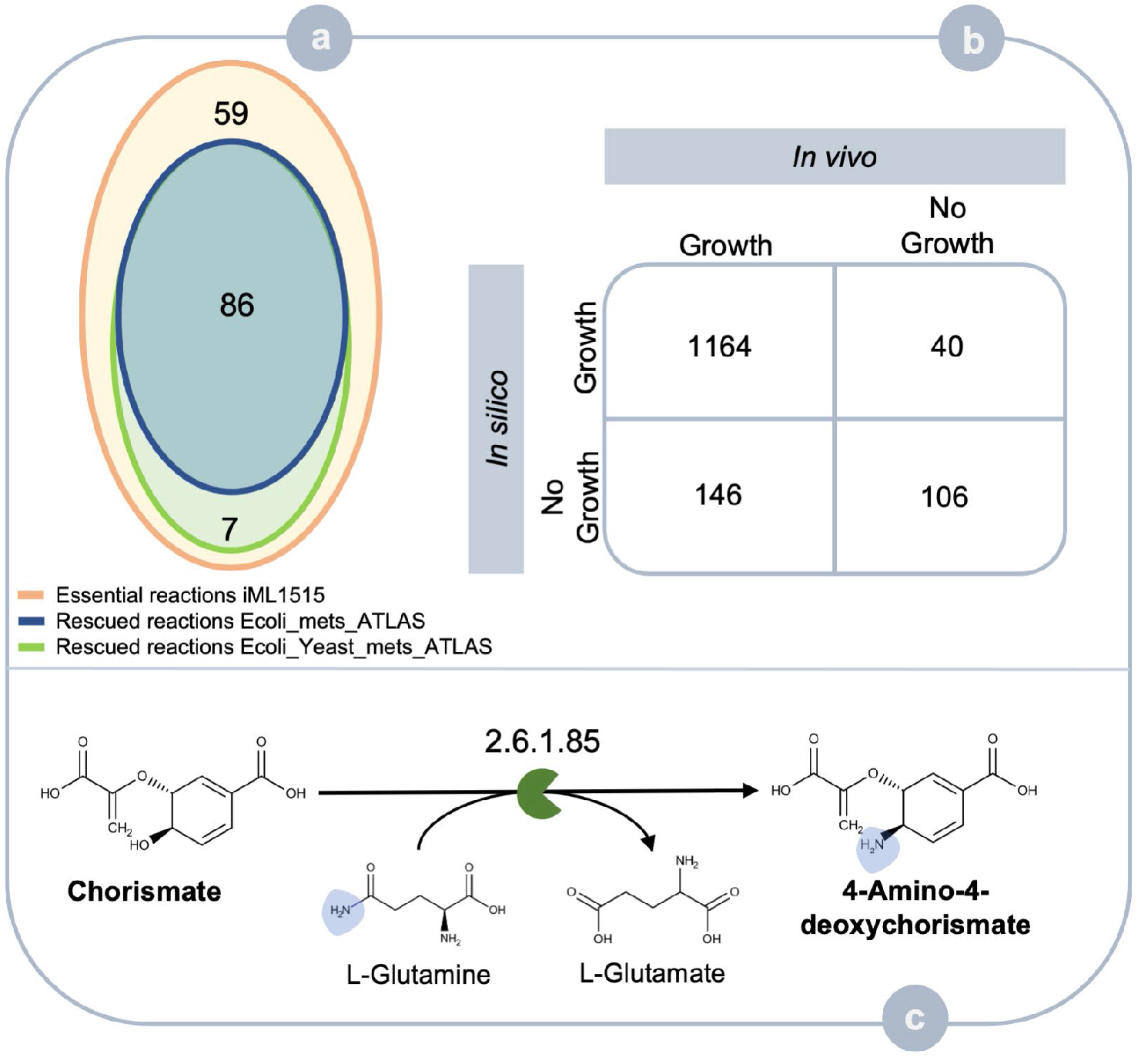
**(A) Comparison of essentiality between iML1515 and the extended networks**. The 146 false negative gene essentiality predictions are linked to 152 (orange) reactions in the model. Using the Ecoli_mets_ATLAS as a reaction pool for gap-filling 86 of these gaps could be characterized (blue) while 92 reactions could be rescued when the Ecoli_Yeast_mets_ATLAS was used as a reaction pool (green). **(B) Contingency matrix for gene essentiality prediction accuracy of iML1515**. The accuracy of the model is equal to 0.872 and MCC equal to 0.488. **(C) Remaining gaps**. The subsets of ATLAS used in this study could not rescue 59 falsely negative reactions. The enzyme 4-amino-4-deoxychorismate synthase (EC 2.6.1.85) remains falsely negative.

### 2.3 Known and novel biotransformations among E. coli metabolites to reconcile model predictions with experimental evidence

The first subset, Ecoli_mets_ATLAS DB, allowed us to suggest at least one thermodynamically feasible solution set for 86 out of 152 false negative cases (Supplementary Table S2). As an example, the enzyme Adenosylmethionine-8-amino-7-oxononanoate transaminase (EC 2.6.1.62) catalyzes the production of 7,8-Diaminononanoate, a precursor of Biotin, from 8-Amino-7-oxononanoate and then encoding gene, bioA, is not essential *in vivo* but it is essential *in silico*. Our approach suggested 116 thermodynamically feasible solution sets that can serve as alternatives to achieve the biosynthesis of Biotin. Figure 4A depicts the original reaction and two alternative reaction sets. In the first alternative, a single reaction can fill the gap. The reaction is novel and follows the same mechanism (EC 2.6.1.-) as the original reaction, however in this case L-Ornithine serves as the donor of the amino-group and L-Glutamate 5-semialdehyde is the byproduct of the reaction. The reaction does not affect the predicted growth rate, does not require any additional enzymatic capability and it improves the overall accuracy of the model with respect to gene essentiality prediction. BridgIT could identify 12 candidate genes with adequate BridgIT score to regulate this reaction. The second alternative requires the addition of two novel reactions to fill the gap; the first reaction converts 8-Amino-7-oxononanoate to 7,8-Diaminononanoate by transferring the amino group from L-Cysteine (EC 2.6.1.-), while Mercaptopyruvate is produced. BridgIT suggests 19 candidate genes. The second reaction is required to balance the production of Mercaptopyruvate by converting it into Hydroxypyruvate (EC 3.3.1.-), following a reaction mechanism that is not part of the original network. However, we could identify one putative sequence to catalyze this reaction.

**Figure 4.**
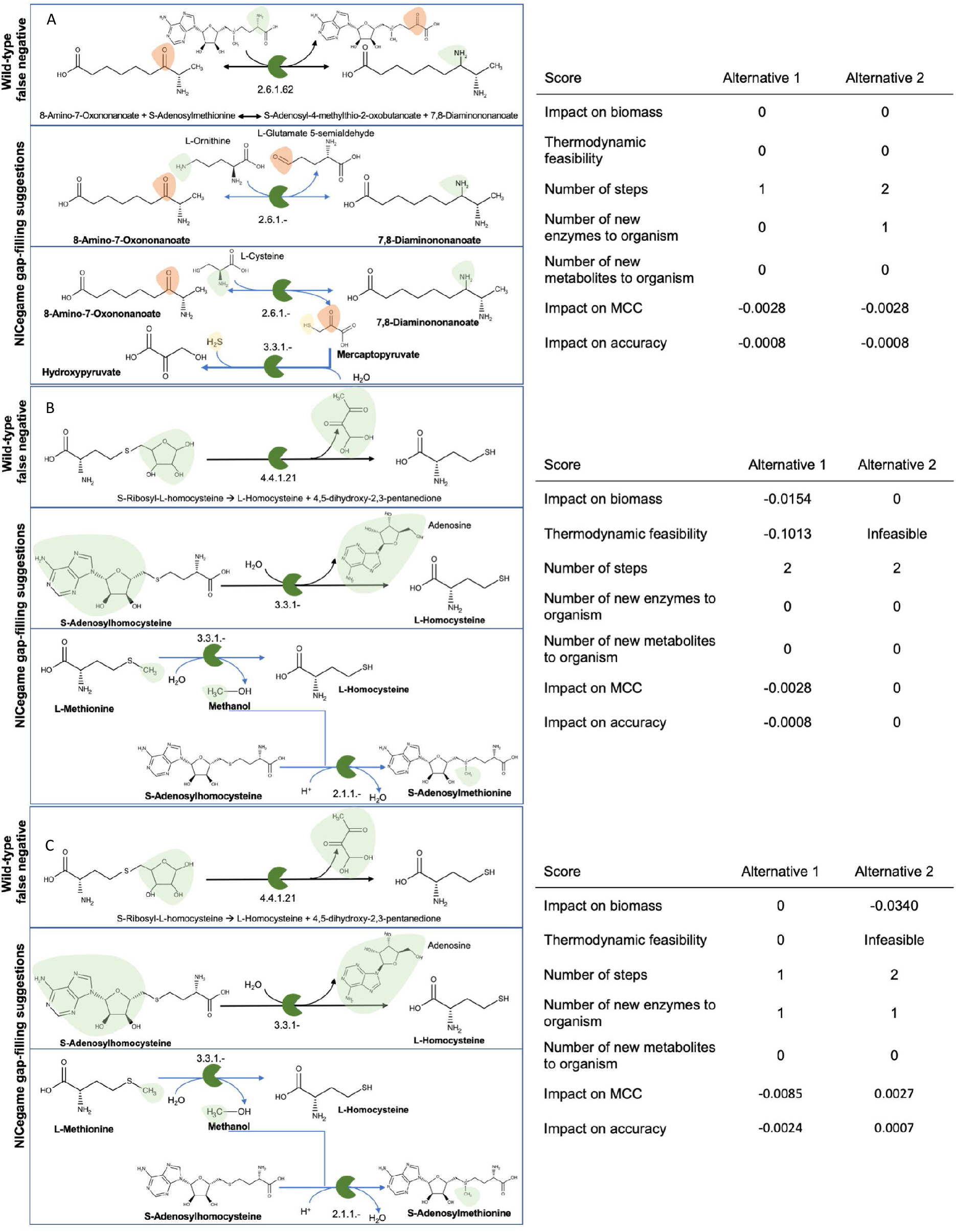
Cases of incorrectly predicted as essential reactions and alternative gap-filling reactions identified using the Ecoli_mets_ATLAS DB. **(A)** The reaction regulated by bioC in the original network, two gap-filling solutions and their scores. **(B)** The reactions catalyzed by luxS in iML1515, one thermodynamically favorable and one thermodynamically infeasible solution. **(C)** The false negative reaction linked with the gene panB and two gap-filling solutions with their scores.

The enzyme 3-methyl-2-oxobutanoate hydroxymethyltransferase catalyzes the production of 2-Dehydropantoate, a precursor of Coenzyme A, from 3-Methyl-2-oxobutanoate (EC 2.1.2.11). This enzyme is the product of the falsely negative gene panB (b0134). The heuristic suggests 29 thermodynamically feasible solution sets. All of the solutions involve the production 2-Dehydropantoate from 3-Methyl-2-oxobutanoate and Formaldehyde (EC 4.1.2.-), which is the orphan KEGG reaction R01216. Our method suggests 26 candidate genes to encode for this biotransformation. Figure 4B depicts the original reaction and two alternative reaction sets. In the first alternative, the side reaction is novel and describes the reduction of Formate to Formaldehyde (EC 1.2.1.-). BridgIT could not identify any candidate sequence to encode for this reaction. In the second alternative, Formaldehyde is produced from 3-Hydroxypropanoate through an acyltransferase (EC 2.3.3.-). However, this novel reaction is not thermodynamically feasible in the desired directionality.

The gene luxS (b2687) is another case of false negative that can be reconciled by gap-filling with the Ecoli_mets_ATLAS DB. The S-ribosylhomocysteine cleavage enzyme, encoded by b2687, is responsible for the production of L-Homocysteine, a precursor of L-Methionine, from S-Ribosyl-L-homocysteine (EC 4.4.1.21). Our workflow suggests 11 thermodynamically feasible solution sets. The first alternative, depicted in Figure 4C, suggests that L-Homocysteine is the product of an amylase acting on S-adenosylhomocysteine (EC 3.3.1.-), that is the KEGG reaction R00192. This enzymatic capability is not part of the original network, however, using BridgIT we could identify one candidate gene to encode for this enzyme. The second alternative requires the same reaction mechanism to produce L-Homocysteine from L-Methionine and a second reaction to balance the bioproduct, Methane. This solution set is thermodynamically infeasible since these reactions form a cycle, L-Methionine → L-Homocysteine → L-Methionine. According to the first law of thermodynamics, the overall thermodynamic driving force through these cycles must be zero, meaning that no net flux is possible through these cycles, making the system infeasible.

### 2.4 Biotransformations among E. coli and yeast metabolites allows the reconciliation of more gaps

A gap-filling reaction pool of biotransformations between *E. coli* and yeast metabolites suggests more alternative solution sets for the already rescued false negative reactions and suggests biochemistry for seven more false negative cases (Supplementary Table S2). One of the additionally rescued reactions is 3-isopropylmalate dehydratase and it accounts for interconversion of 2-Isopropylmaleate to 3-Carboxy-2-hydroxy-4-methylpentanoate (EC 4.2.1.33). The reaction is part of the Leucine Biosynthesis pathway, and it is regulated by two genes, leuD (b0071) and leuC (b0072). Figure 5A shows the original reaction and two alternative pathways. The first set of reactions describes the production of 4-Methyl-2-oxopentanoate, a precursor of Leucine, from Butanoyl-CoA via a 4-step path. Three of the reactions are novel and BridgIT can identify candidate genes for two of them, while the solution includes also the KEGG reaction R01176. The second set of reactions accounts for the synthesis of 4-Methyl-2-oxopentanoate again from Butanoyl-CoA, via a 3-step path of novel reactions, however, it is thermodynamically unfavorable. Both solution sets involve the metabolite 3-Methylbutanal that is not part of the original iML1515 metabolic network. The compound has been characterized^16^ as an alternative substrate of the enzyme 3-hydroxypropionaldehyde dehydrogenase (b1300) but has not been detected as part of the metabolome of *E. coli* to our knowledge.

**Figure 5.**
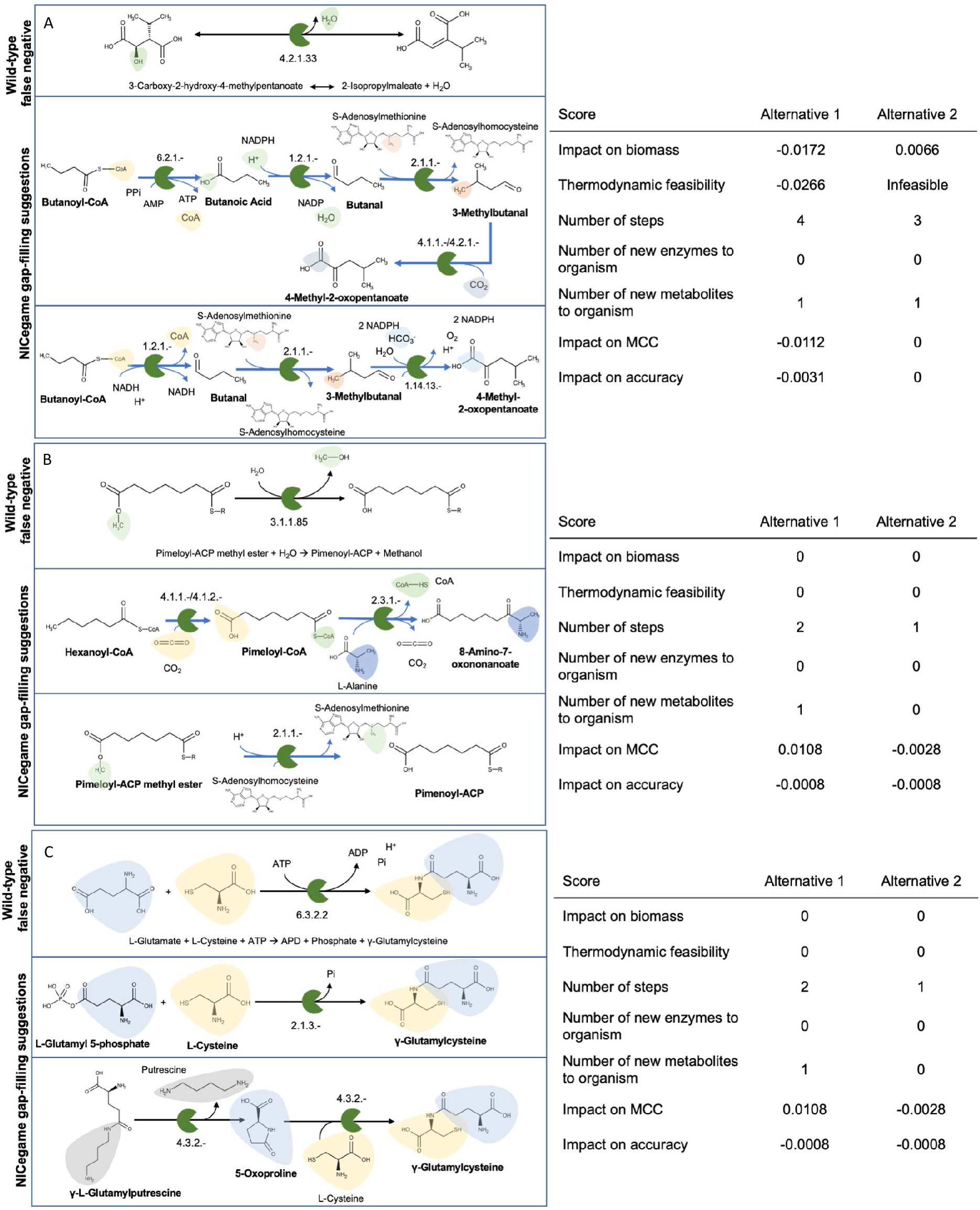
Cases of incorrectly predicted as essential reactions and alternative gap-filling reactions identified using the Ecoli_Yeast_mets_ATLAS DB. **(A)** The false negative reaction regulated by leuCD in the original network, one thermodynamically favorable and one thermodynamically infeasible gap-filling solution and their scores. **(B)** The reactions catalyzed by bioH in iML1515, one lowly ranked and one highly ranked gap-filling solution. **(C)** The false negative reaction linked with the gene gshA and two gap-filling solutions with their scores. The first alternative is generated by the Ecoli_mets_ATLAS DB whereas the second one is generated by the Ecoli_Yeast_mets_ATLAS DB.

Pimeloyl-ACP methyl ester esterase (EC 3.1.1.85) is an enzyme encoded by the false negative gene bioH (b3412). The enzyme is part of the Biotin Biosynthesis pathway, and it is responsible for the hydrolysis of Pimeloyl-ACP methyl ester to Pimeloyl-ACP, a precursor of Biotin. This reaction is rescued by the Ecoli_mets_ATLAS DB, which provides three alternative solution sets of one reaction each to fill this gap. One of them, (Figure 5B, alternative 2), suggests that Pimeloyl-ACP is produced by the transfer of the methyl group from Pimeloyl-ACP methyl ester to S-Adenosylhomocysteine forming S-Adenosylmethionine (EC 2.1.1.-). BridgIT provides 11 genes to regulate this novel reaction. The Ecoli_Yeast_mets_ATLAS DB provides one additional solution set, (Figure 5B, alternative 1), of two steps, a novel reaction, and the KEGG reaction R03210, describing the synthesis of 8-Amino-7-oxononanoate, a successor metabolite of Pimeloyl-ACP in the Biotin Biosynthesis pathway, from Hexanoyl-CoA and Pimeloyl-CoA as an intermediate metabolite. Pimeloyl-CoA is not part of the original reconstruction but has been recently shown^17^ that it can serve as the acyl chain donor of the 8-amino-7-oxononanoate synthase (bioF). BridgIT suggests 21 candidate genes to encode for this function with bioF being among them. In the original reconstruction bioF, which uses Pimenoyl-ACP as substrate, is essential but the gene is not essential *in vivo*. Interestingly, this solution set provides an alternative precursor for Biotin and thus can resolve the false negative case of bioC (b0777) that is responsible for the synthesis of Malonyl-CoA methyl ester, a precursor of Pimenoyl-ACP and thus Biotin. However, this solution is rejected since it adds redundancy to the model, having an MCC (Matthews Correlation Coefficient) score equal to 0.0108 since the genes fabZ (b0180) and fabH (b1091) and are falsely positive after the addition of this solution set to the network.

The gene gshA (b2688), which regulates the synthesis of γ-Glutamylcysteine from L-Cysteine and L-Glutamate (EC 6.3.2.2), is not essential *in vivo* but it is essential *in silico*. This gap can be resolved by the Ecoli_mets_ATLAS DB. The heuristic provides 12 alternative thermodynamically feasible solution sets to fill in this gap. The solution that is ranked higher according to our criteria is depicted in Figure 5C. γ-Glutamylcysteine is produced by L-Cysteine and L-Glutamyl 5-phosphate, while Orthophosphate is released (EC 2.1.3.-). The Ecoli_Yeast_mets_ATLAS DB provides 3 additional thermodynamically favorable solution sets. All the 3 solutions involve the metabolite 5-Oxoproline as an intermediate. 5-Oxoproline is not part of the metabolome of the original reconstruction, it was, however, has been detected^18^ in *E. coli*. The most well performing solution is shown in Figure 5C. The first reaction, which describes the degradation of γ-L-Glutamylputrescine to Putrescine and 5-Oxoproline, is novel, whereas the second reaction is a KEGG reaction (R02743) reconstructed in ATLAS. BridgIT identifies the gene chaC as a candidate to encode for this metabolic function.

### 2.5 Gene annotation of metabolic gaps identifies new functions in E. coli

The heuristic suggests over 7,000 reactions, known and novel, to gap-fill the metabolic network of *E. coli*, and over 6,600 among them are part of thermodynamically feasible solution sets. We used BridgIT to identify candidate sequences in the genome of *E. coli* to catalyze these reactions. 6,319 reactions had an adequate BridgIT score (see Methods) and are assigned to 2,165 EC numbers. Finally, we suggest 590 candidate promiscuous genes in the genome of *E. coli* to catalyze 6,118 reactions. An example is shown in Figure 6. BridgIT could assign adequate similarity scores between the ATLAS novel reaction and five KEGG reference reactions and identifies the gene chaC as promising to catalyze this novel reaction.

**Figure 6.**
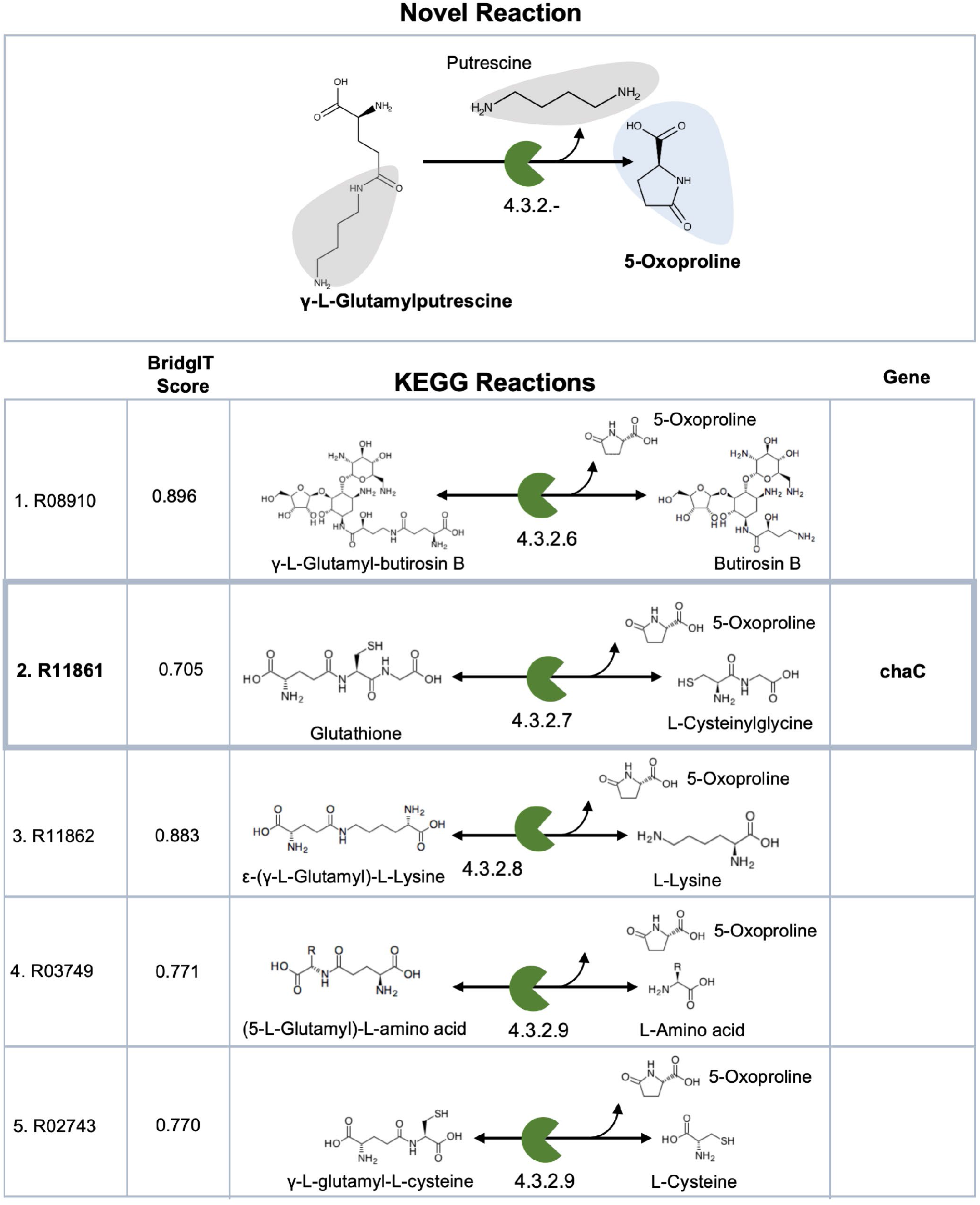
Suggesting catalyzing genes using BridgIT. BridgIT could identify 5 reactions that are similar to the novel reaction that accounts for the degradation of γ-L-Glutamylputrescine to 5-Oxoproline and Putrescine. Out of these five reactions only R11861 is linked to a sequence in the genome of *E. coli*.

### 2.6 An updated genome-scale model and database of E. coli metabolism shows increase essentiality prediction accuracy

Our approach allows to expand the original metabolic network by 77 reactions and 9 metabolites. We suggest 35 genes to associate with these 77 reactions, 2 of them are not part of the original reconstruction (Supplementary Table S3). These 35 genes include only the top-rated hits provided by BridgIT. Using the criteria and the ranking method mentioned above we extracted an updated version of *E. coli’s* strain MG1655 metabolism, iEcoMG1655 (Figure 7). The updated reconstruction includes 2,450 network reactions, 1,176 metabolites, and 1,517 genes, while it has an enhanced accuracy in gene essentiality prediction. iEcoMG1655 achieves a MCC equal to 0.6025 and an ACC (accuracy) equal to 0.9190, while iML1515’s performance is 0.4879 and 0.8722, respectively, for the conditions examined in this study.

**Figure 7.**
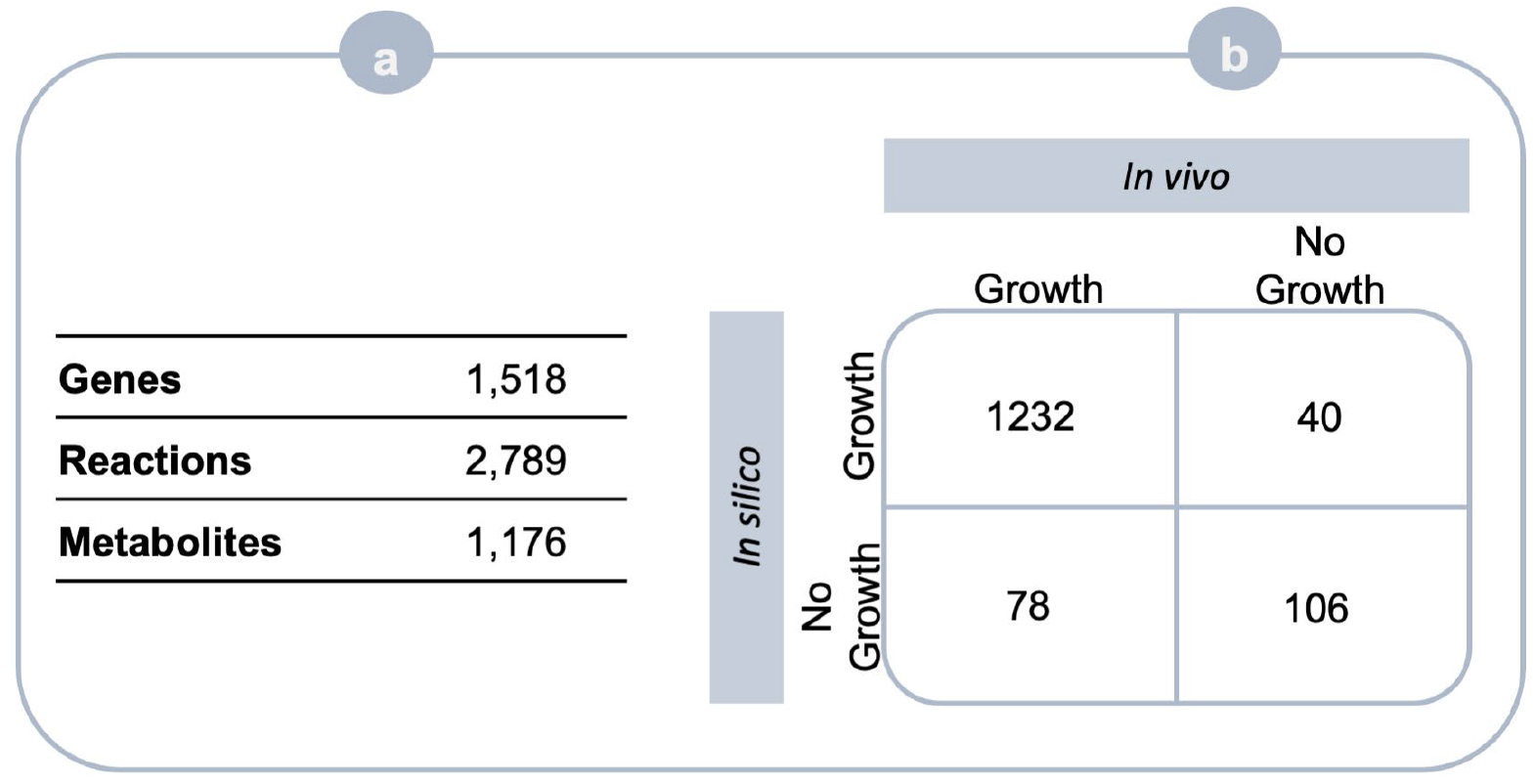
**(A) The iEcoMG1655 network statistics**. The gap-filled network contains 77 novel reactions, two additional genes and nine new metabolites. **(B) Contingency matrix for gene essentiality prediction accuracy of iEcoMG1655**. Our approach could reconcile 58 out of the 146 FN gene essentiality predictions leading to an increased accuracy, i.e., MCC = 0.6025.

ArcA and LacA are the 2 added genes. ArcA (b4401) is part of the ArcAB (aerobic respiratory control) regulatory system^19^. ArcA and ArcB have been shown to regulate the expression of oxygen-requiring pathways^19^. ArcAB has been also known to participate in the proper expression of catabolic genes for pyruvate utilization and sugar fermentation pathways^19^. In the expanded reconstruction the gene regulates the hydrolysis of N2-succinyl-L-arginine to Urea and N2-Succinyl-L-ornithine (EC 3.5.3.-), providing an alternative pathway to compensate for the knockout of argG (b3172).

LacA (b0342) encodes the enzyme Galactoside O-acetyltransferase that catalyzes the transfer of an acetyl group from acetyl-CoA to the 6-hydroxyl of some galactopyranosides^20^. The enzyme is known to act on a broad range of substrates and can acetylate galactosides, thiogalactosides, glucosides, and lactosides^20^. In the expanded reconstruction it is part of the Lipopolysaccharide Biosynthesis / Recycling and catalyzes the degradation of Dodecanoyl-KDO2-lipid IV(A) to KDO2-lipid A, compensating for the knockout of lpxM (b1855).

The 33 genes that already are part of the model show substrate or mechanism promiscuity. In the original reconstruction galK (b0757) encodes for the enzyme Galactokinase, responsible for the phosphorylation of D-Galactose (EC 2.7.1.6). galK shows substrate promiscuity since BridgIT suggests it can phosphorylate ADP-D-glycero-D-manno-heptose, 5-Methylthio-D-ribose, D-glycero-beta-D-manno-Heptose 7-phosphate, 6-(Hydroxymethyl)-7,8-dihydropterin, and D-Ribose 5-phosphate. On the other hand, xapA (b2407), acts as a glycosyltransferase (EC 2.4.2.-) in the original reconstruction whereas BridgIT suggests it can encode for phosphotransferases (EC 2.7.4.-) and ydfG (b1539), acts as dehydrogenase (EC 1.1.1.-) in the iML1515 network, but we suggest it can act as a carbon-carbon lyase (EC 4.1.1.-, 4.1.2.-).

Although NICEgame suggests thermodynamically feasible alternatives to rescue 92 reactions, solutions were not added in the updated reconstruction for 14 reactions for two reasons. The first reason not to include a gap-filling solution in the updated reconstruction is that all the solution sets identified by our approach add redundancy in the metabolic network, resulting in an increase of the false positive gene essentiality predictions. The second reason not to include any solution set for a rescued reaction was that we had an indication that an essential or a falsely negative gene catalyzes the suggested biochemistry.

The newly included biochemistry allows the reconciliation of 78 false negative essential reactions and 68 false negative essential gene cases. The biochemistry added to form the iEcoMG1655 is the top-rated among the alternatives, based on the knowledge that it is available to this day. Our approach allows users to revisit and reevaluate the suggested solution sets for each rescued reaction, in case new quantitative and qualitative data are released, by suggesting alternative solution sets.

The amended metabolic network iEcoMG1655 can serve as a tool to explore the undiscovered metabolic capabilities of *E. coli*. The library of alternative gap-filling solutions can be used to postulate experimentally testable hypotheses to shed light on the underground metabolism of *E. coli*. Our method and the library of alternative solution sets can also be used as a recourse in metabolic engineering, to design strains with improved performance, i.e., higher biomass or product yield. Our workflow is applicable to any genome-scale metabolic model of prokaryotic organisms. NICEgame allows the characterization and curation of metabolic gaps at the reaction and enzyme level leading towards fully annotated genomes.

### 2.7 The NICEgame workflow offers improved gap-filling performance

To evaluate the performance of NICEgame against existing gap-filling approaches we performed three comparative studies. In the first study, we repeated the generation of gap-filling alternative solutions using our in-house algorithm but only known biochemical reactions, i.e., the Ecoli_Yeast_mets_KEGG DB, a subset of the KEGG database, as a pool for the gap-filling. The second study compares our gap-fling algorithm to published algorithms, i.e., the algorithms included the RAVEN and COBRA toolboxes, using the Ecoli_Yeast_mets_ATLAS DB. Last, we compared NICEgame against the CarveMe gap-filling approach.

The Ecoli_Yeast_mets_KEGG DB allows the identification of thermodynamically feasible gap-filling solutions for 53 out of the 152 target reactions (Supplementary Table S4), contrary to 93 rescued false negative reactions by the Ecoli_Yeast_mets_ATLAS DB. The average number of solutions per rescued reaction is 2.3 for the Ecoli_Yeast_mets_KEGG DB and 252.2 for the Ecoli_Yeast_mets_ATLAS DB. However, the Ecoli_Yeast_mets_KEGG DB suggests solutions for 8 reactions that the Ecoli_Yeast_mets_ATLAS DB cannot rescue. Further analysis of the KEGG reactions that can substitute for the 8 *E. coli* reactions was performed to understand why the ATLAS database could not capture these solutions (Supplementary Table S5).

The gap-filling approach implemented in the RAVEN toolbox also uses a MILP (Mixed-Integer Linear Programming) algorithm. We used the input parameters so that is the problem is to include as few reactions as possible from the database in order to satisfy the model constraints, i.e., the mass balances and the basal growth rate under the defined media. The problem is very similar to the one defined by the NICEgame, however the RAVEN approach does not account for the generation of alternative solutions. This results in obtaining thermodynamically feasible solutions for 67 out of the 152 target reactions (Supplementary Table S6). The gap-filling package that is included in the COBRA toolbox is also a MILP algorithm, that minimizes the number of added reactions from the database to the model, in order to achieve a given metabolic task. The COBRA toolbox algorithm gives different weights to the reactions of the database, penalizing the uptakes and transporters in comparison to metabolic reactions. The algorithm can account for alternative solutions, by assigning a bigger weight to reactions that have already appeared in previous solutions. To test the method, we demanded 10 alternatives per target reaction. However, the same solution can reappear and there is no systematic enumeration of all minimal solution sets. (Supplementary Table S7). The reaction ANPRT is such an example. The algorithm identifies the ATLAS reaction ‘rat45874’ as a solution four times, alternatives 1, 2,7, and 10. The size of the remaining six solutions ranges between two and three reactions per solution set.

The CarveMe gap-filling approach is another gap-filling method based on a MILP formulation, that aims to add the minimal number of reactions from the Universal Bacterial model^21^ to the genome-scale model under curation. The method does not generate alternative solutions sets. The usage of a different database leads to the identification of different gap-filling alternatives. For example, the Ecoli_Yeast_mets_ATLAS DB can gap-fill for the target reaction AICART with solution sets of one reaction, whereas the Universal Bacterial model provides a solution of 20 reactions. Using the CarveMe gap-filling we identified thermodynamically feasible solutions for 33 target reactions and reconcile 24 gaps that are not curated by the NICEgame. However, the method suggests transporter, e.g., CAt6, CITt13, Cuabc, and pseudo-reactions, e.g., sink_4hba_c, as part of the solutions.

Overall, NICEgame outcompetes existing gap-filling approaches since it achieves an exhaustive and systematic enumeration of gap-filling solutions of the minimal or bigger size. Using the ATLAS of Biochemistry as a reaction pool for gap-filling allows the reconciliation of more gaps. The thermodynamic evaluation of the alternatives in combination with our scoring system allows us to choose biologically relevant solutions. The integration of the BridgIT tool in the NICEgame workflow offers the users an inclusive genome-scale model gap-filling, from the metabolite to the enzyme level.

## 3. Methods

In this study, we present our gap-filling approach, which comprises seven main steps. The workflow produces a merged metabolic network by connecting the metabolism of the organism with a reaction database, i.e., the ATLAS of Biochemistry. Then, it attempts to create a functional network by substituting essential reactions, to the original metabolic network, for biomass production with reactions coming from the reaction database. Candidate annotated and non-annotated genes in the organism that can catalyze the suggested biochemistry are identified. The alternative solutions are then evaluated and ranked.

### 3.1 Reconciliation of annotation

The suggested workflow was implemented on the most recently published *E. coli* model, iML1515^2^. The model is derived from 1,515 genes, associated with 2,266 reactions. It integrates 1,569 metabolic reactions and 1,169 unique metabolites across two compartments, the cytosol and the periplasm, and the extracellular space. Since the ATLAS of Biochemistry is KEGG^22^-based, both compound and reaction IDs of the wild-type *E. coli* model needed to be translated to KEGG notation. The whole process required manual search in BiGG^23^ and KEGG databases. The most suitable KEGG ID was matched to the metabolites in the model, based on the BiGG ID and the name of each metabolite. Every metabolite must have its unique ID so that the stoichiometric matrix of the model is properly generated. Thus, in order to avoid conflicts, in the case of different compounds with the same KEGG ID, such as lipids and stereoisomers, the KEGG ID was assigned only once. 909 out of the 1,169 metabolites were mapped to a KEGG ID. Apart from the KEGG database, the metabolites of the iML1515 were also mapped to the SEED^24^ database, so that thermodynamic constraints can be imposed on the model. 1,106 out of the 1,169 metabolites are mapped to a unique SEED ID.

### 3.2 Databases used for gap-filling

In this project, we examined the performance of the ATLAS of Biochemistry as a reaction pool for gap-filling. For this project, the updated version of ATLAS^14^ was used. It includes 10,935 KEGG metabolites integrated into 149,052 novel and already known enzymatic reactions. 5,764 of the ATLAS reactions are exact reconstructions of KEGG reactions.

Due to the vast amount of information integrated into ATLAS, in this project we used two subsets of ATLAS. More specifically, the Ecoli_mets_ATLAS DB is a subset of the ATLAS database that contains only reactions that integrate compounds, from the intracellular and the extracellular space of the cell, that are already part of the iML1515 genome-scale model. To this end, the ATLAS database was converted in a pseudo-GEM format and any reactions that integrated compounds that do not belong in the *E. coli* metabolic network were removed. We thus examined whether the gaps in the model can be reconciled by expanding only the reaction space while not increasing the metabolite space. Likewise, the the Ecoli_Yeast_mets_ATLAS DB contains only ATLAS reactions that integrate compounds that are already part of the iML1515 and the Yeast8^3^ genome-scale models. In this case, more information was extracted from ATLAS in a controlled way, expanding both the reaction and metabolite space of the original metabolic network. Likewise, the Ecoli_Yeast_mets_KEGG DB contains only KEGG (2018 version) reactions among metabolites included in the iML1515 and the Yeast8^3^ genome-scale models. The metabolic network of yeast was chosen since *E. coli* is often cultivated in yeast extract, and it is thus probable that parts of the missing metabolome exist in the yeast metabolic reconstruction. For the gap-filling with the CarveMe approach, the universal bacterial model^21^, a compartmentalized model contains transporters and pseudo-reactions, was used as a database. The size of the databases is shown in Table 1.

**Table 1:** Size of the databases that were used for gap-filling.

|  | Reactions | Metabolites |
| --- | --- | --- |
| Ecoli_mets_ATLAS | 9,810 | 778 |
| Ecoli_Yeast_mets_ATLAS | 13,298 | 1,050 |
| Ecoli_Yeast_mets_KEGG | 1,756 | 1,128 |
| Universal Bacterial model | 5,532 | 2,861 |

### 3.3 Gap-filling formulation

The gap-filling algorithm generates binary use variables for each reaction in the database. These variables indicate whether flux is allowed through a reaction or not. The gap-filling algorithm is in reality a parsimonious algorithm whose objective is to minimize the number of active reactions in the metabolic network, demanding at the same time a basal flux through the biomass reaction in the wild-type model. The mathematical formulation of the MILP problem is:

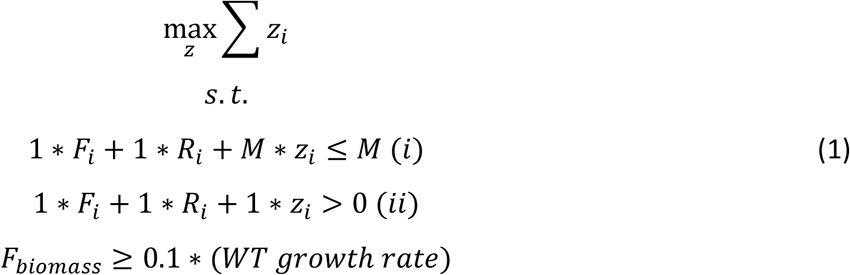

where Fi stands for the flux variables of the irreversible forward reactions, Ri are the flux variables of the irreversible backward component reactions of the reversible reaction i, Fbiomass is the flux variable of the irreversible forward biomass reaction, WT growth rate is the growth rate of the wild-type model, M is a big-M value, m is a small value and zi are the binary use variables.

Every time the solver identifies a solution, the solution is integrated as a cut constraint to the MILP problem, so the solver cannot identify the same solution more than once:

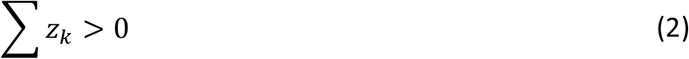

We generated solutions of the minimum and the subsequent size. To avoid the generation of long pathways we demanded that the minimum solution size is less than 10 and the subsequent solution can be at most 5 reactions bigger than the minimum size solution.

### 3.4 Identification of metabolic gaps

Gene essentiality data^25^ were used to identify putative false negative reactions. We considered M9 glucose minimal media and aerobic conditions and the wildtype biomass reaction as an objective function. We performed a single gene deletion analysis, where a gene was considered essential *in silico* if the growth rate of the knock-out mutant was less than 10 % of the growth rate of the wildtype. This analysis revealed 258 genes essential *in silico* with 105 of them being essential in vivo, while 7 of them are not part of the experimental data. We identified all reactions associated with the 146 remaining genes, 200 in total, and after a single reaction deletion analysis, we concluded that 152 of them are essential in silico. We consider that these 152 are falsely essential and thus constitute the target reactions for gap-filling.

### 3.5 Scoring the alternatives

The output of the framework, for each gap-filled model, is a set of ranked alternatives for each rescued reaction. The main criteria for ranking the different alternatives are the thermodynamic feasibility of the solution, which means the system cannot violate the second law of thermodynamics, and minimum impact on the model, that means that the more a solution alters the biochemistry and the predictive capability of the model the lower it is ranked.

Thermodynamics-based Flux Balance Analysis (TFA)^26^ is carried out for each alternative in order to examine the maximum biomass yield under thermodynamic constraints. In order to examine the maximum biomass yield for each alternative the rescued reaction is blocked, and flux is allowed through a set of the reactions of the alternative. Then a TFA is carried out. These values are compared with the performance of the wildtype GEM and the ratios of the wildtype GEM to the gap-filled GEM are calculated. Then, 1 is subtracted from the ratio of the maximum biomass yield of the two models. If the result of the subtraction is greater than 0, the addition of the alternative to the GEM leads to a lower performance compared to the original model, whereas, in the case that the result of the subtraction is less than 0, the addition of the alternative to the GEM leads to higher performance compared to the original model. If a gap-filled GEM does not predict growth when it is thermodynamically restricted, the alternative is rejected. The performance of the gap-filled models without thermodynamic constraints is also tested. FBA is carried out for each alternative in order to examine the maximum biomass yield. The results are analyzed similarly to the TFA test.

The number of reactions of each alternative is also tested. Since the set of reactions of each alternative replaces one reaction in the model, 1 is subtracted from the number of reactions in the solution set. Since usually organisms favor shorter paths, the alternatives that integrate fewer reactions are ranked higher than those that integrate more reactions.

Furthermore, the metabolites integrated into each reaction are examined. For every unique non-native metabolite, 1 point is added. An extra point is added for every reaction that is linked to a 3rd level EC number that is not included in the original GEM. The integration of such reactions also entails the integration of new enzymatic capabilities into the model.

Lastly, we test the ability of the models to properly predict gene essentiality. To this end, the overall accuracy (ACC) and Matthews Correlation Coefficient (MCC) are calculated for each gap-filled model and are compared to the WT.

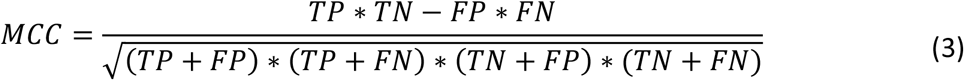

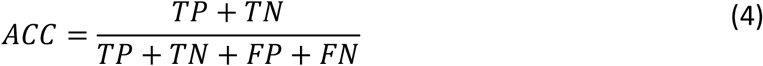

where TP stands for True Positive, TN for True Negative, FP for False Positive, and FN false negative gene essentiality model predictions.

The values of all the scores are added, and the alternatives are ranked. The closer the absolute value of the score is to 0, the more similar the performance and the biochemistry of the gap-filled model is to the original model.

### 3.6 Enzyme annotation with BridgIT method

In this study, for the annotation of ATLAS reactions, we used the online version of the BridgIT method with default parameters as discussed in the original paper^15^. BridgIT method is inspired by the theory of lock and key, assuming two similar reactions will be catalyzed by the same enzyme. More precisely, BridgIT takes into account the reactive site and its neighborhood in similarity calculations, since the rest of the structure doesn’t interact with the enzymatic binding pocket.

BridgIT method compares the similarity of each input reaction with all the non-orphan characterized metabolic reactions cataloged in the KEGG database (reference reactions) and proposes the enzymes associated with the most similar reference reactions as the best candidate for the input reaction. Therefore, BridgIT systematically screens for the best promiscuous candidate enzymes that might be able to catalyze the input reaction. The degree of similarity or probability of catalyzation is quantified in the BridgIT score, ranging from 0 (no similarity) to 1 (identical). The optimal threshold value for the BridgIT score is 0.3, meaning predictions with a score higher than 0.3 are considered promising^15^.

The output of BridgIT for each input reaction is a list of ranked enzymes ordered descending based on BridgIT score along with their EC number. Then, EC number is used to query the Uniport^27^ database for the corresponding protein sequences in the organism of interest, in this study Escherichia coli K12. Finally, the BridgIT output annotated with protein sequences in Escherichia coli K12 is used for gap-filling.

## Software

This work was supported by EPFL through the use of the facilities of its Scientific IT and Application Support Center. We performed gap-filling using the defined MILP formulation in MATLAB (2016a and 2018a) and IBM ILOG Cplex 12.7.1 as a solver. The simulations were run on a High-Performance Computing Cluster of 408 nodes. We used 2 CPUs per simulation and 3875 MB per CPU. One simulation was defined for a unique combination of parameters. The analysis of the gap-filling solutions was performed on mac info in MATLAB 2017a and IBM ILOG Cplex 12.7.1 as a solver. The gap-filling with RAVEN was performed with Gurobi Optimizer Version 9.3 as a solver. The gap-filling with COBRA and CarveMe approaches were performed in python 3.6 and IBM ILOG Cplex 12.8.0 as a solver.

## Acknowledgments

Funding for this work was provided by Swiss National Science Foundation (SNSF): grant 200021_188623, NCCR Microbiome grant 51NF40_180575, SystemsX.ch MicroScapeX grant 2013/158, and SystemsX.ch MalarX grant 2013/155, European Union’s Horizon 2020 research and innovation programme grants: PacMen, under the Marie Sklodowska-Curie grant agreement No 72228, and ShikiFactory100, under grant agreement 814408, Swedish Research Council Vetenskapsradet (grant no. 2016-06160), and the École Polytechnique Fédérale de Lausanne.

## Author Contributions

EV, ACP and VH conceptualized the study. EV, ACP, HM, YF, NH, MA, JH performed the data curation. EV, ACP and YF developed the software. EV, ACP, MA and VH developed the methodology. EV, ACP, and HM carried out the formal analysis. Investigation was conducted by EV, ACP, HM, YF and VH. EV and ACP wrote the manuscript. EV, ACP, and VH developed the visualizations. All authors edited and reviewed the manuscript. ACP, SP and VH managed and supervised the project. VH acquired the funding and the resources.

## Corresponding author and material availability

Further information and requests for resources should be directed to and will be fulfilled by the corresponding author, Vassily Hatzimanikatis. The workflow is available at the LCSB GitHub https://github.com/EPFL-LCSB/NICEgame. The code for TFA is available at https://github.com/EPFL-LCSB/mattfa. The ATLAS of Biochemistry is available at the LCSB database https://lcsb-databases.epfl.ch/pathways/atlas/ and the BridgIT tool at https://lcsbdatabases.epfl.ch/pathways/Bridgit.

## Declaration of interests

The authors declare no competing interests.

## Supplementary Table title and legends

**Supplementary Table S1**.

False negative gene essentiality model predictions and the reactions that constitute the targets of the gap-filling algorithm.

**Supplementary Table S2**.

All gap-filling solutions identified by the heuristic along with the corresponding scores with the Ecoli _mets_ATLAS and the Ecoli_Yeast_mets_ATLAS databases.

**Supplementary Table S3**.

The top-rated solutions and the suggested catalyzing enzymes used to update the reconstructions.

**Supplementary Table S4**.

All gap-filling solutions identified by the heuristic with the Ecoli_Yeast_mets_KEGG DB.

**Supplementary Table S5**.

Rescued reactions by the Ecoli_Yeast_mets_KEGG DB and not the Ecoli_Yeast_mets_ATLAS DB.

**Supplementary Table S6**.

All gap-filling solutions identified by the RAVEN gap-filling approach.

**Supplementary Table S7**.

All gap-filling solutions identified by the COBRA gap-filling approach.

**Supplementary Table S8**.

All gap-filling solutions identified by the CarveMe gap-filling approach.

## Notes

https://github.com/EPFL-LCSB/NICEgame

### Competing Interest Statement

The authors have declared no competing interest.

### Summary of Updates

We capitalized the first letter in the first name of the first author.

